# Individual mouse mitotic chromosomes exhibit cell type-specific differences in biomechanical properties

**DOI:** 10.64898/2026.09.07.749823

**Authors:** Karen E. Brown, Shubhadeep Patra, Andrew Dimond, Korak Kumar Ray, Sherry Cheriyamkunnel, Bhavik Patel, Do Hyeon Gim, Chad Whilding, Marta Llobet Ayala, Mehmet Efe Kilic, David S. Rueda, Amanda G. Fisher

## Abstract

Cyclic episodes of chromosome compaction and de-condensation are features of eukaryotic cell division that aid mitotic segregation and help prevent aneuploidy. While biophysical data on mitotic chromosome structure has been previously obtained, heterogeneity within samples can confound analyses and precludes direct like-for-like comparisons. To circumvent this, we employed advanced flow cytometry to purify specific metaphase chromosomes with biotinylated telomeres from stably engineered mouse cells. We show that ESC-derived metaphase chromosomes 3 and 19 display distinct properties but share a conserved force-dependent mechanical response. In contrast, chromosome equivalents isolated from NSCs and preB cells show markedly different force-dependent responses, reflecting progressive differentiation stages. Covalent crosslinking of ESC-derived chromosomes alters biomechanical properties to mimic equivalents from more differentiated cells. Collectively, these results highlight the need to isolate specific, homogeneous metaphase chromosome samples to accurately decipher their complex behaviours.

## Introduction

Chromosomes are major carriers of genetic information that is passed between generations through reproduction, via meiosis, and through organismal development, via mitosis. There is huge interest in understanding the biophysical properties and structure of chromosomes, particularly during mitosis, when wholesale compaction of genetic material occurs that is critical for achieving correct segregation between daughter cells^1–5^. This is particularly challenging because chromosomes are complex polymers composed of DNA, RNA, histones, and a plethora of other chromatin and structural components, and because their assembly radically changes once cells enter mitosis^6–9^. Attempts to measure and model mitotic chromosome behaviour span more than 40 years of research and have included diverse experiments on amphibian, insect, and mammalian cells from which chromosomes have been extracted, manipulated *in vivo* or *ex vivo*, and stretched^10–14^. Pioneering early experiments using newt and human chromosomes suggested a near-linear relationship between stretching force and extension for up to 5 times their original length, with mitotic chromosomes returning to a native state upon release. This reversible spring-like behaviour was not preserved with much larger chromosome extensions^11^, was eroded by multiple successive extensions^12^ and probably does not adequately explain the response of mitotic chromosomes to much slower rates of extension^15^.

The recent advances in optical tweezer approaches that enable high-resolution force measurements together with fluorescence visualisation of metaphase chromosomes has opened the door to new studies of mitotic chromosome structure and biomechanics^15–19^. In their landmark paper^16^, Wuite and Hickson showed that metaphase chromosomes, derived from a BirA-TRF1-expressing human osteosarcoma cell line U2OS, showed non-linear stiffening with increasing mechanical load. Since this behaviour was distinct from that predicted by classical polymer models, a hierarchical worm-like chain model has been proposed where metaphase chromosomes are viewed as a heterogeneous assembly of non-linear chains. A potential limitation of this interesting study was that it was not possible to determine which of the many different human chromosomes had contributed to the analyses. This could be important since human chromosomes differ widely in size, gene density, and shape, as well as possessing distinctive architectural and epigenetic features^20–22^. To address this potential source of heterogeneity, we have opted for a more precise approach using specific individual mouse metaphase chromosomes that were purified by flow cytometry^23,24^ and subsequently mechanically manipulated using optical tweezers. This, we reasoned, would enable us to accurately assess the biomechanical responses of a homogeneous population of individual chromosomes derived from a non-disease associated and karyotypically stable source. In addition, mouse metaphase chromosomes share a similar acrocentric shape and display broadly similar gene densities (two-fold differences between chromosomes), as compared to human chromosomes, where seven-fold differences are reported^20^.

To isolate native unfixed mouse metaphase chromosomes, we developed an advanced flow cytometry approach in which chromosomes were isolated based on a bivariate karyotype that reflects AT:GC ratio and mass^23,25^. Using this system, we have previously examined the repertoire of proteins that remain chromosome-associated through mitotis^23^ in mouse embryonic stem cells (ESCs), profiled candidates evicted by mitotic kinase-mediated phosphorylation^26^, and determined factors that are retained differentially by active versus inactive X chromosomes in female mitotic cells^27^. Here, we used an analogous approach to examine the biomechanical properties of individually purified mouse chromosomes 3 and 19. To do this, we engineered stable cell lines that constitutively expressed BirA-TRF1 and showed that sustained expression and biotinylation of telomeres did not significantly alter chromosome size or the growth and viability of cells. Force extension data indicated that despite chromosome-specific differences in stiffness, chromosomes 3 and 19 derived from ESCs share a conserved force-dependent mechanical response. However, comparisons of metaphase chromosome 3 isolated from different cell types revealed a trend of altered mechanical behaviours that correlated with differentiation stage. We further show that these trends were consistent with the widespread changes in chromatin architectures that occur when cells differentiate.

## Results

### Engineering mouse ESCs with biotinylated telomeres through stable expression of BirA-TRF1

To study the biomechanical properties of mouse metaphase chromosomes using optical tweezers, we opted for a strategy where the chromosome telomeres could be tethered to optically trapped beads and mechanically manipulated^16^. Cells were engineered in which all chromosome telomeres were biotinylated by the stable introduction of a fusion protein between BirA (biotin ligase) and telomere repeat-binding factor 1 (TRF1)^16,28^. The *BirA-TRF1* construct was inserted into the mouse *Rosa26* locus of mouse ESCs using CRISPR/Cas9- based targeting, as outlined in Figure 1a. Individual ESC clones were molecularly screened for insertion of *BirA-TRF1* in one or both *Rosa26* alleles using PCR and primer pairs that distinguish the wild-type *Rosa26* locus (WT) from the targeted locus (KI) (Figure 1a and 1b). Multiple clones were generated from which a heterozygote KI clone (c10) was identified where streptavidin labelling of all telomeres was evident throughout the cell cycle (Figure S1a), including during mitosis. Figure 1c shows a representative single optical section of BirA-TRF1 c10 ESCs at metaphase, with DAPI (blue) and streptavidin (green) labelling of condensed acrocentric chromosomes and telomeres, respectively, and where close examination of multiple Z-series stacks confirmed streptavidin labelling of all telomeres. To check that the expression of BirA-TRF1 at telomeres did not in itself, alter metaphase chromosome size, we performed DNA FISH using chromosome paints and metaphase spreads prepared from WT and c10 (BirA-TRF1) ESCs. As shown for chromosomes 3 and 19, a pair of signals were detected with these probes in all cells (representative images shown in Figures 1d and S1b). An assessment of the chromosome area labelled by each DNA-FISH paint in WT versus BirA- TRF KI (c10) samples showed that, while chromosome 3 was larger than chromosome 19, there was no significant differences in size between equivalents derived from WT and BirA- TRF1 targeted cells (Figure 1e).

**Figure 1.**
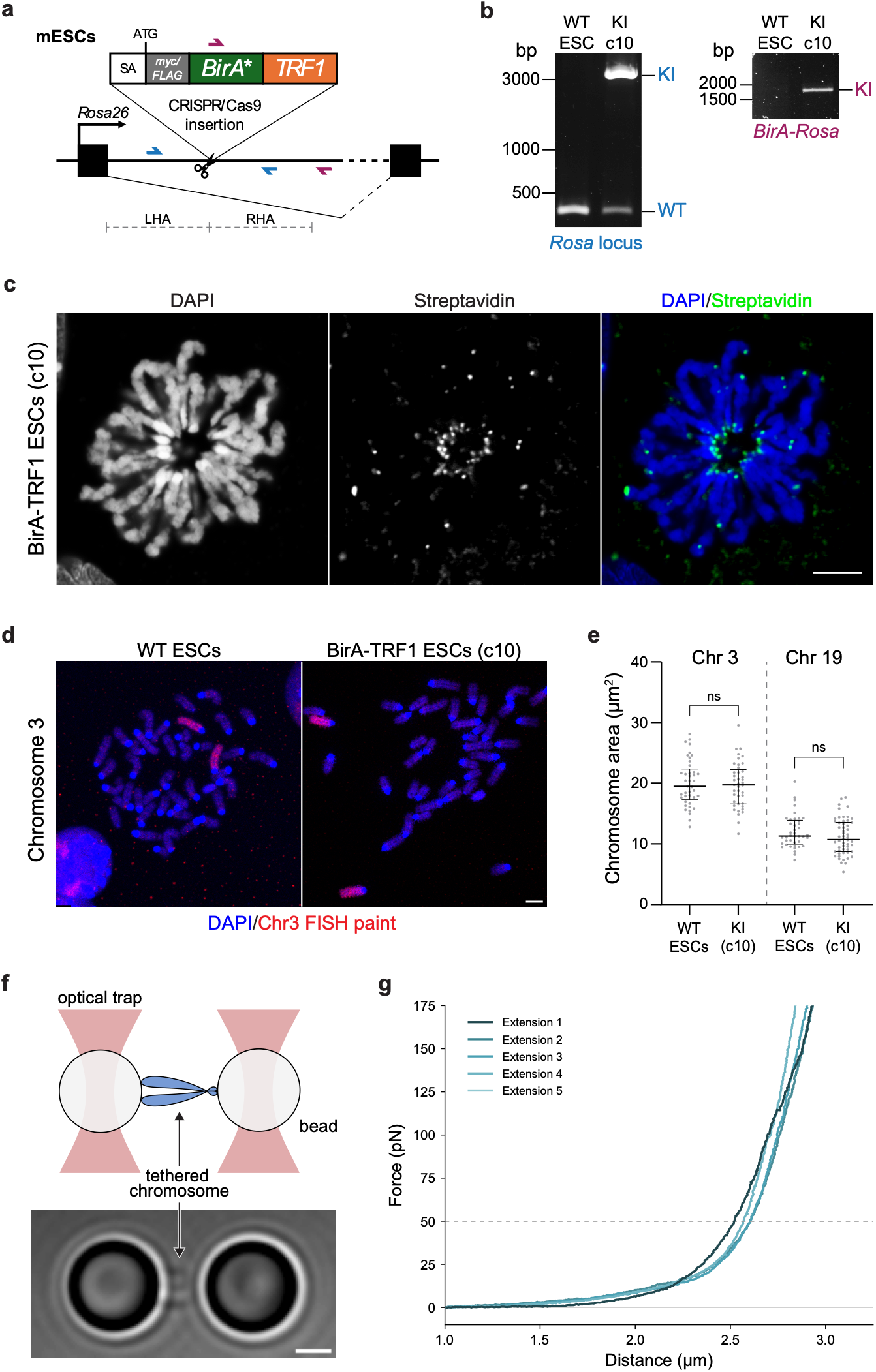
Stable expression of BirA-TRF1 in mouse ESCs for biophysical interrogation of mitotic chromosomes. **a,** Strategy for BirA-TRF1 stable expression in ESCs by CRISPR/Cas9-mediated knock-in (KI) to the *Rosa26* locus, showing splice acceptor (SA); left/right homology arms (LHA/RHA); and primers to verify insertion (red and blue arrows). **b,** PCR confirming heterozygous *BirA-TRF1* KI to the *Rosa26* locus in clone 10 (c10) ESCs using primers shown in (a). Wild-type (WT) and knock-in (KI) alleles are indicated. **c,** Biotin incorporation at telomeres in c10 ESCs detected with Alexa Fluor 488- labelled streptavidin, shown in a single z-slice of a representative metaphase cell; note biotin labelling can be observed at all telomeres across z-slices. Scale bar = 5 µm. **d,** Representative metaphase spreads from parental WT and c10 (BirA-TRF1-expressing) ESCs, hybridised with chromosome 3 paint probes (red). **e,** Size quantification of chromosomes 3 and 19 from WT and c10 ESC metaphase spreads (N = 41, 41 for chr 3, and N = 41, 54 for chr 19), based on DAPI segmentation and identified by chromosome paints. Plots show median and interquartile range (unpaired two-tailed t-tests with Holm-Šídák correction, ns = not significant). **f,** Schematic (top) and brightfield image (bottom) of a metaphase chromosome tethered between two optically trapped beads. Scale bar = 2 µm. **g,** FECs for repeated extensions of a single representative mouse ESC chromosome. The corresponding relaxations are not shown for clarity.

Finally, we confirmed that chromosomes derived from BirA-TRF1c10 (c10) could be captured and manipulated using optically trapped streptavidin-coated beads. The brightfield image shown in Figure 1f (lower image) shows an individual mouse acrocentric metaphase chromosome. Once captured, the individual chromosomes were successively extended and relaxed multiple times (Figure 1g). Consistent with previous studies using BirA-TRF1 labelled human metaphase chromosomes^16^, mouse metaphase chromosomes could be pulled to very high forces, up to hundreds of pN. Successive force-extension curves (FECs) for the same chromosome remained broadly similar (Figure 1g). However, we observed a large heterogeneity among unsorted mouse chromosome FECs (Figures S1c and S1d) and the corresponding distribution of chromosome extensions at a reference force of 50 pN (Figure S1e). This is similar to what had been previously observed for human chromosomes^16^ derived from an unsorted mixture of all chromosomes. As a result, it was important to isolate specific individual chromosomes to distinguish heterogeneity in behaviours beyond that arising from sample heterogeneity.

### Metaphase chromosomes exhibit conserved force-dependent biomechanics despite chromosome-specific variations

We used advanced flow cytometry to prospectively identify and purify individual subsets of metaphase chromosomes^23^. Previously, we have used this technique to comprehensively catalogue ‘mitotic bookmarking proteins’ that co-purify with native metaphase chromosomes^23^, to dissect how alterations in H3K9me3 heterochromatin alters this repertoire^24^, and to identify a cadre of proteins that selectively associate with either the active or inactive X chromosome in mitosis^27^. Here, we used the same experimental pipeline to separately purify chromosomes 3 and 19 from BirA-TRF1 (c10) mouse ESCs. Briefly, cells were arrested with demecolcine; native metaphase chromosomes were released and stained with DNA dyes chromomycin A3 and Hoechst 33258 to generate a bivariate karyotype where chromosomes could be easily discriminated (Figure 2a, left, Figures S2a-b), purified, and captured on streptavidin beads (Figure 2a, right). DNA FISH analysis of flow sorted samples of metaphase chromosome 3 or 19 (Figures 2b and S2b) confirmed that >90% and >92% of chromosomes within these samples labelled with 3-specific or 19-specific DNA paints, respectively.

**Figure 2.**
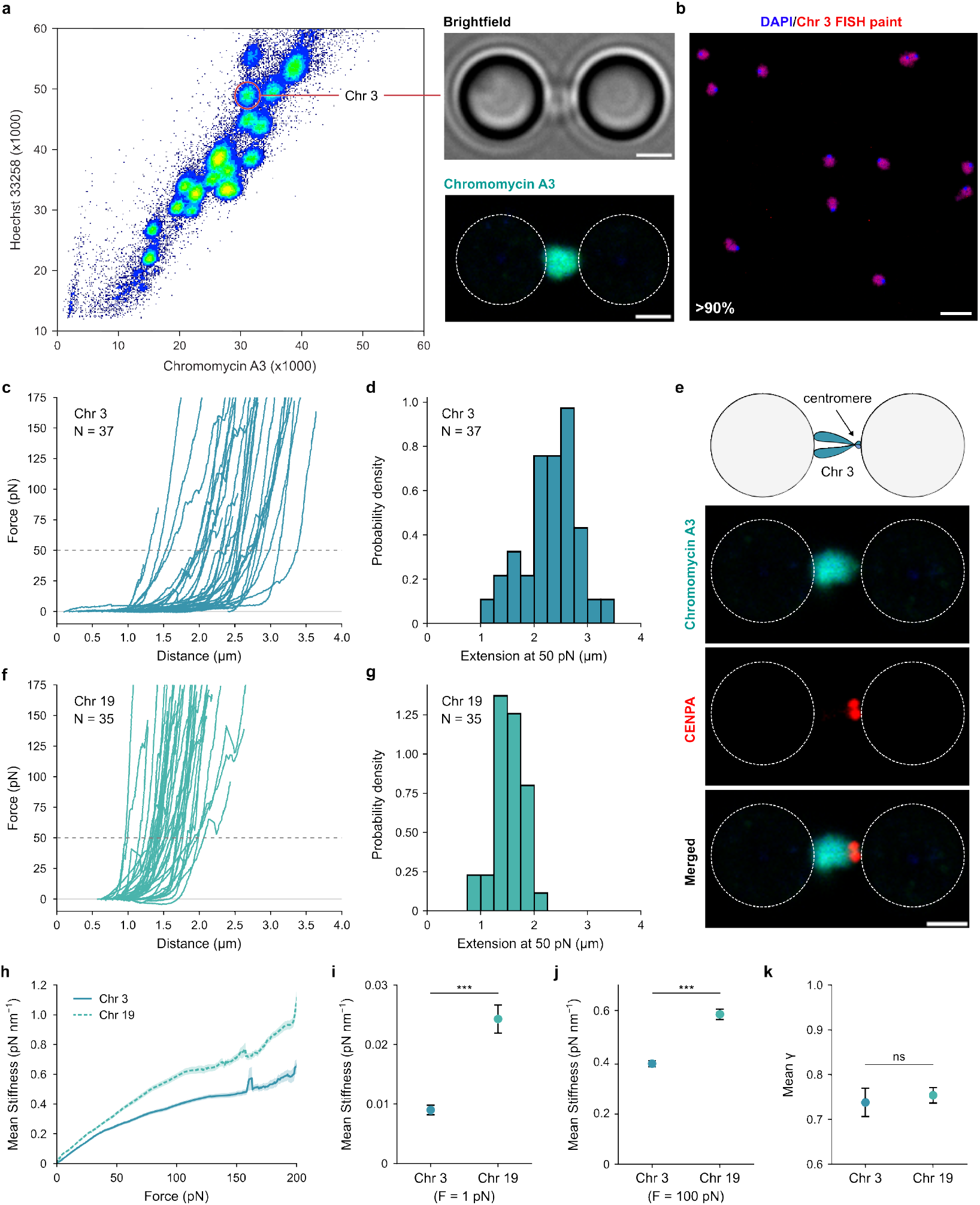
Biophysical properties of individual ESC mitotic chromosomes isolated by flow cytometry. **a,** Representative flow cytometry visualisation (left) of stained mitotic chromosomes (Hoechst 33258/chromomycin A) from c10 ESCs. Gating for purification of chromosome 3 is shown. Representative brightfield and fluorescence images (right) of a single tethered ESC chromosome 3. Scale bars = 2 µm. **b,** Purity of flow cytometry-purified chromosome 3 confirmed as >90% by hybridisation with chromosome 3 paint probes (N = 345), visualised with DAPI (blue) and paint (red). Scale bar = 10 µm. **c,** FECs for flow-purified ESC chromosome 3, showing the first extensions (N = 37). **d,** Histogram of ESC chromosome 3 extensions at 50 pN (N = 37). **e,** Schematic of tethered mouse chromosome 3 with asymmetric centromere position (upper), compared to representative fluorescence images (lower) of a tethered flow-purified ESC chromosome 3, stained with CENPA antibody (red) to mark the centromere. Scale bar = 2 µm. CENPA staining adjacent to one bead, indicative of short-to- long arm attachment, was observed in all cases (N = 8/8). **f,** FECs for flow-purified ESC chromosome 19, showing the first extensions (N = 35). **g,** Histogram of ESC chromosome 19 extensions at 50 pN (N = 35) **h,** Force-stiffness curves for ESC chromosomes 3 (N = 37, n = 123) and 19 (N = 35, n = 114). Mean stiffness ± s.e.m. (shading) is shown. **i–k,** Mean stiffness at 1 pN (i) and 100 pN (j), and mean *γ* (k) for ESC chromosomes 3 and 19 (Welch’s t-test, *** = p < 0.001, ns = not significant). Error bars correspond to s.e.m. Extensions were calculated from the first FEC of each chromosome, while stiffness and *γ* were calculated from all FECs.

Analysis of the FECs of metaphase chromosome 3 showed a heterogeneous range of chromosome behaviour (Figures 2c and S2c). We observed a wide distribution of extensions at 50 pN (Figure 2d), with a mean extension of 2.30 ± 0.08 µm, where the error represents the standard error of the mean (s.e.m). Importantly, this is similar to the mean of the unsorted mixture of ESC metaphase chromosomes (2.2 ± 0.1 µm; Figure S2d). While the variation in extensions for the subset of chromosome 3 was expectedly smaller than the one for the unsorted chromosome population, the difference was not statistically significant (Figure S2e). Thus, our results demonstrated that even a purified sample of a specific chromosome displays a range of variability.

One explanation for this remaining heterogeneity might be that it is due to a different tethering geometry of the chromosomes on the beads—for example, if they are tethered by the long chromosome arms versus the short arms, or if the tethering includes one or both acrocentric arms. To examine this possibility, we labelled chromosome 3 with an antibody to CENPA, marking the centromeres and allowing us to discern the spatial orientation of the tethered chromosomes. Visualisation of the residual chromomycin A3 (green) alongside CENPA-labelling (red) showed a near-identical orientation in 8 of 8 chromosome 3 examples that were examined (example shown in Figure 2e). This suggests that the major chromosome orientation captured in our experiments arose from an initial capture of the two long arms (or alternatively the two short arms) on one streptavidin-coated bead (outlined in white, left in images shown), followed by tethering of the remaining two arms by the other bead upon stretching by buffer flow (to the right in the images shown). This suggests that the heterogenous chromosome behaviour we observed could not be solely attributed to differences in tether orientation.

To examine whether a smaller chromosome behaves similarly, we repeated this analysis using purified metaphase chromosomes 19 from c10 ESCs as an exemplar. Due to the smaller size of chromosome 19, we had to employ slightly smaller beads to enable capture (2.17 µm vs 4.35 µm). As shown in Figures 2f and 2g (and Figures S2c and S2d), FECs of the individual metaphase chromosome 19 confirmed the reduced size of chromosome 19 as compared to chromosome 3, with a mean extension length at 50 pN of 1.54 ± 0.04 µm. The corresponding distribution of extensions for chromosome 19 showed significantly less variation than the unsorted population of chromosomes, but was not significantly lower than for chromosome 3 (Figure S2e). This suggests that mouse metaphase chromosomes show considerable heterogeneity regardless of the size of the specific chromosome.

To quantitatively compare the specific biomechanics of chromosomes 3 and 19, we calculated the stiffness of these chromosomes as a function of the applied force. Previous studies had observed that human metaphase chromosomes become stiffer (*i.e.*, more resistant to extension by force) with increasing force^16^. We observed a similar force-dependent increase in stiffness for both chromosomes 3 and 19 (Figures 2h and S2f). Remarkably, however, we found that mouse chromosome 19 was stiffer than mouse chromosome 3. This difference in stiffness between the two chromosomes persisted over the entire range of applied forces, from a very low force (1 pN), before the chromosomes undergo appreciable extension, to a high force (*e.g.*, 100 pN), where the chromosomes are under significant tension (Figures 2i and 2j, respectively). Interestingly, the stiffnesses at low forces for both mouse chromosomes were nearly an order of magnitude lower than that reported for human chromosomes^16^. While the cause for this is not clear, we note that this difference is not due to our specific purification process, since the stiffness at 1 pN for the mixed pool of unsorted and unstained mouse chromosomes was similarly low (0.011 ± 0.001 pN nm^−1^).

Previous studies^16^ have modelled the force-dependence of chromosome biomechanics with the power-law relation, *K* ∝ *F^γ^*, where the exponent %, called the stiffening exponent, describes how the chromosome stiffness (*K*) changes with the applied force (*F*). A larger value of *γ* indicates that stiffness increases more rapidly with applied force, whereas a smaller exponent reflects a more gradual increase in resistance to extension. We found that a single non-zero value of *γ* failed to capture the force-dependence of stiffness for mouse metaphase chromosomes. As a result, we elected to calculate a local value of *γ* for each point of the FECs (see Methods) (Figure 2k). For chromosome 3, our calculations yielded a mean *γ* value of 0.74 ± 0.03, in line with the value reported previously for human metaphase chromosomes^16^. For chromosome 19, the mean *γ* was 0.75 ± 0.02, nearly identical to chromosome 3 (Figure 2k). Therefore, the biomechanics for both chromosomes exhibit a similar force-dependence, despite chromosome-specific differences in stiffness. The conserved force-dependent biomechanical response of chromosomes 3 and 19 suggests a common chromosomal architecture between the two, one that is likely shared by all chromosomes in mouse ESCs (see Discussion).

### The biomechanical properties of mouse metaphase chromosomes change with differentiation stage

To examine how the biomechanical properties of individual metaphase chromosomes compare to equivalent chromosomes isolated from other cell types, we differentiated c10 ESCs into neural stem cells (NSCs; Figure 3a, left). This was achieved by withdrawing LIF, culturing the resulting neural progenitors with specialised N2B27 media and EGF/FGF2 growth factors and isolating self-renewing NSCs grown on laminin (c10 NSCs)^29^. As anticipated these cells lost expression of several pluripotency markers including *Oct4* and *Nanog* (as compared to undifferentiated c10 or WT mouse ESCs) but had increased *Nestin* and *Pax6* gene expression (Figure 3a, right). C10-derived NSCs also displayed changes in morphology, with loss of Oct4 protein expression and increased Nestin expression (Figure S3a) consistent with acquisition of stable neural identity. Biotinylated mouse metaphase chromosome 3 was isolated from these NSCs (c10) by flow cytometry (Figure 3b) for biomechanical analysis with optical tweezers. DNA FISH analysis of the sorted chromosome 3 from c10 NSCs confirmed >92% purity of these preparations. As shown in Figure 3c (and Figure S3b), the FECs of mouse chromosome 3 isolated from neural progenitors (c10 NSCs) were broadly similar to equivalents isolated from ESCs (c10). However, these chromosomes were more compact, with a mean extension at 50 pN of 2.16 ± 0.06 µm (Figure 3d).

**Figure 3.**
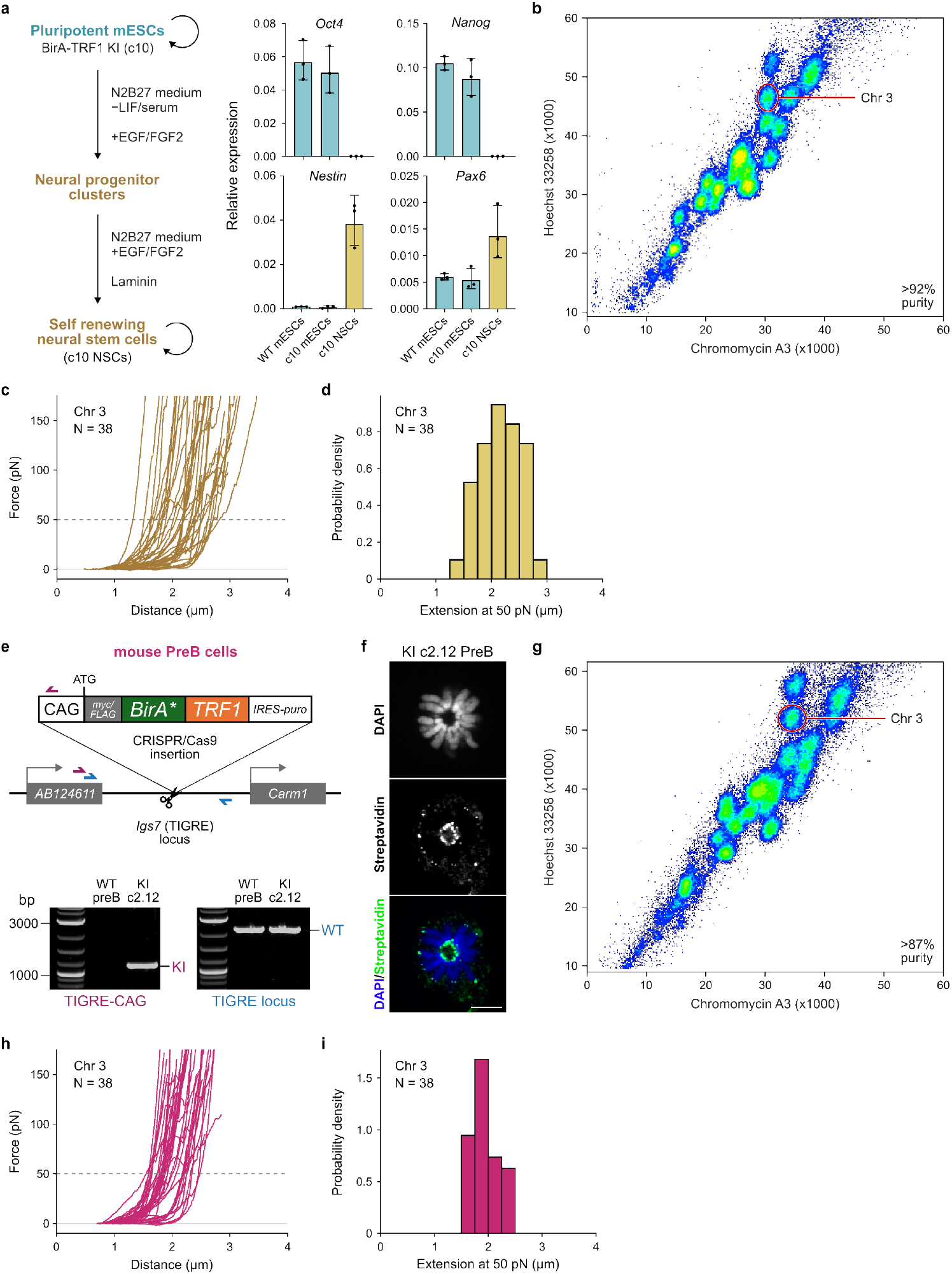
Properties of mitotic chromosome 3 from mouse NSCs and preB cells. **a,** Derivation of BirA-TRF1-expressing NSCs from c10 ESCs (left). Differentiation-induced loss of pluripotency and gain of NSC-markers verified by qRT-PCR (right), with gene expression (geometric mean) in parental WT and BirA-TRF1 KI c10 ESCs (blue) and c10-derived NSCs (brown) normalised to *β-actin* and *Gapdh*. Error bars correspond to s.d. **b,** Representative flow karyotype of stained mitotic chromosomes from c10 NSCs, with gating of chromosome 3 and percentage purity from a typical experiment, determined by hybridisation to chromosome 3 paint probes (N = 189). **c,** FECs for flow-purified NSC chromosome 3, showing first extensions (N = 38). **d,** Histogram of c10 NSC chromosome 3 extensions at 50 pN (N = 38). **e,** Generation of BirA-TRF1-expressing mouse preB cells by CRISPR/Cas9-mediated knock-in to the *TIGRE* locus (top; CAG = CAG promoter). PCR confirming heterozygous *BirA-TRF1* integration in clone 2.12 (c2.12) preB cells (below), where WT and KI alleles are detected. **f,** Biotin incorporation at telomeres in BirA-TRF1-expressing c2.12 preB cells, detected with Alexa Fluor 488-labelled streptavidin, shown in a single z-slice of a representative metaphase cell; note biotin labelling is observed at all telomeres across z-slices. Scale bar = 5 µm. **g,** Representative flow karyotype of stained mitotic chromosomes from c2.12 preB cells, with gating of chromosome 3 and percentage purity from a typical experiment, determined by hybridisation to chromosome 3 paint probes (N = 388). **h,** FECs for flow-purified preB chromosome 3, showing first extensions (N = 38). **i,** Histogram of preB chromosome 3 extensions at 50 pN (N = 38). Extensions were calculated from first FECs for each chromosome.

To compare the properties of metaphase chromosome 3 equivalents between pluripotent or multipotent stem cells with those derived from a self-renewing source of differentiated cells, we selected mouse preB cell lines and biotinylated telomeres by stably introducing *BirA-TRF1* into the mouse *Igs7*/TIGRE locus using CRISPR/Cas9-mediated insertion as outlined in Figure 3e. In this instance, puromycin selection (IRES-puro) was included to allow selection of BirA-TRF1 KI cells. We identified a heterozygous clone (KI c2.12 preB) carrying a wildtype *Igs7*/TIGRE allele and a KI allele (primer pairs and molecular analysis used to confirm this are provided in lower panels of Figure 3e). Immunofluorescent staining of KI c2.12 preB cells with streptavidin (green) confirmed biotinylation at telomeres of mouse acrocentric chromosomes in this cell line, as illustrated in a representative metaphase image provided in Figure 3f. To isolate metaphase chromosomes, KI c2.12 preB cells were treated with demecolcine as described previously and released chromosomes stained with Hoechst 33258 and chromomycin A3 prior to flow sorting (Figure 3g). Sorted samples were typically ∼90% pure as validated using mouse chromosome 3-specific paint and DNA FISH. The FECs of individual metaphase 3 chromosomes (Figures 3h and S3b) showed that these chromosomes were substantially more compact than equivalents isolated from either ESCs or NSCs, with mean extension at 50 pN of 1.94 ± 0.04 µm (Figure 3i).

These data indicate that genetically identical chromosomes from different cell types acquire different biomechanical properties as they develop. As these characteristics are heritable, and not the result of changes in DNA sequence, it is tempting to speculate that they have an epigenetic origin. To better understand these differences, and the underlying trends, we directly compared the FECs of chromosome 3 from mouse ESC, NSC, and preB cells. As noted above, the mean extensions of chromosome 3 at 50 pN is progressively smaller from ESCs to NSCs to preB cells (Figures 4a and S4a). This decrease was also correlated with a reduction in the variation within the corresponding extension distributions (Figure S4b). The ESC chromosome extension distribution was the most heterogeneous, while the distribution for preB chromosomes was significantly more tightly clustered.

**Figure 4.**
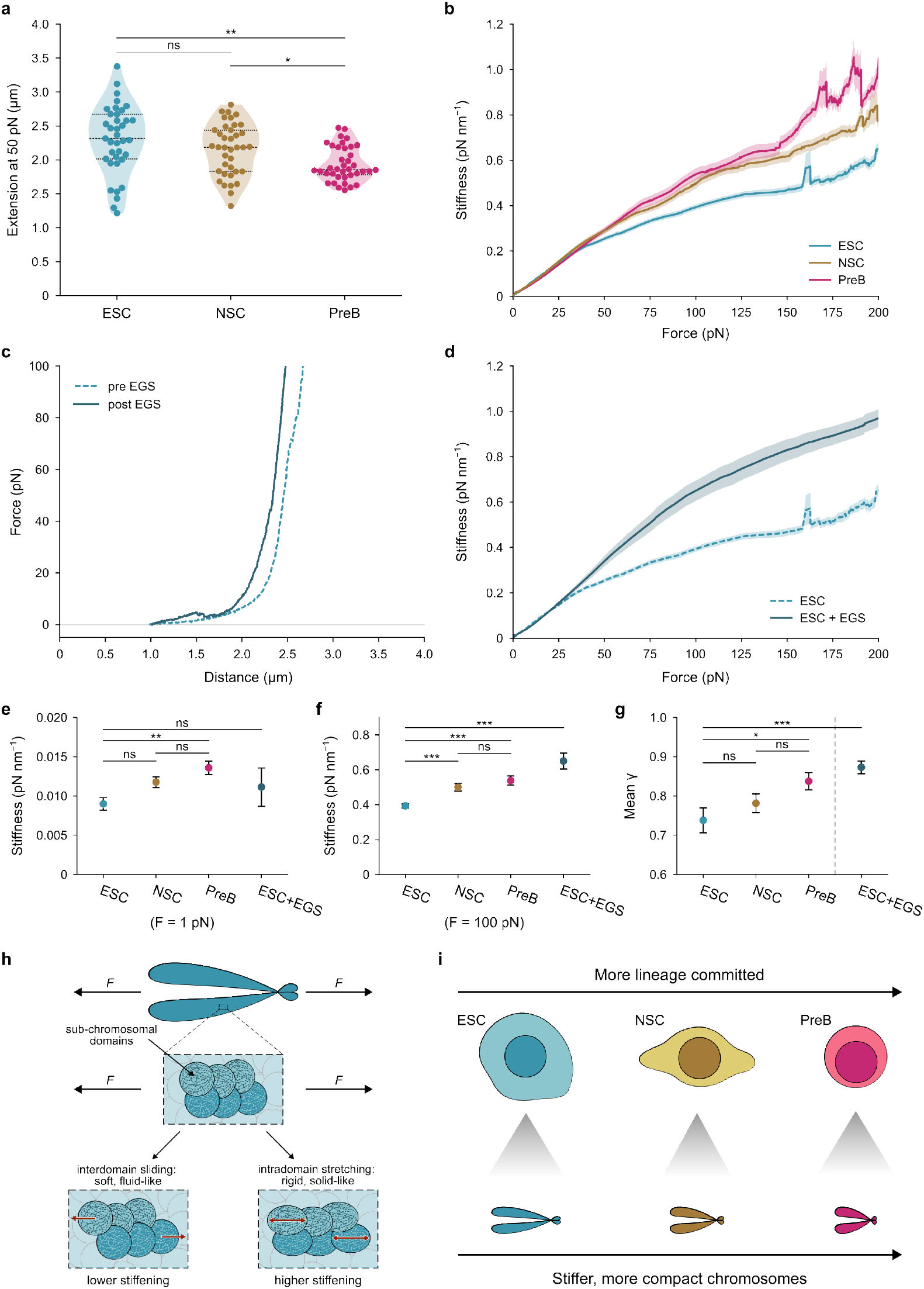
Cell-type specific biophysical properties of mitotic chromosomes. **a,** Violin plots of extensions of flow-purified mitotic chromosome 3 from ESCs (N = 37), NSCs (N = 38), and preB cells (N = 38) at 50 pN, showing median and quartiles **b,** Force-stiffness curves for chromosome 3 from ESCs (n = 123), NSCs (n = 124), and preB cells (n = 132). Mean stiffness ± s.e.m (shading) is shown. **c,** Representative FECs for a single ESC chromosome 3, before (dotted line) and after (solid line) crosslinking with EGS. **d,** Force-stiffness curves for EGS-crosslinked ESC chromosome 3 (solid line, N = 26, n = 119), compared to untreated ESC chromosome 3 (dotted line, as shown in b). Mean stiffness ± s.e.m is shown. **e–g,** Mean stiffness at 1 pN (e) and 100 pN (f), and mean *γ* (g) for chromosome 3 from ESCs (n = 123), NSCs (n = 124), and preB cells (n = 132), as well as EGS-crosslinked chromosome 3 (n = 119) from ESCs **h,** Schematic showing how increased chromatin crosslinking may alter the biomechanical behaviour of metaphase chromosomes in response to force through changes in subchromosomal architecture. **i,** Summary of the changes in biomechanical properties of mitotic chromosomes in progressively more differentiated cell types. (Welch’s t-tests with Holm-Šídák correction, ns = not significant, * = p_adj_ < 0.05, ** = p_adj_ < 0.01, *** = p_adj_ < 0.001).

We calculated chromosome stiffness as a function of force for both NSC- and preB- derived metaphase chromosomes. The force-dependent stiffnesses for NSC- and preB- derived chromosomes 3 were markedly different from the identical ESC-derived chromosome, with higher values corresponding to the most differentiated cells (Figure 4b). These changes in mechanical properties of mouse chromosome 3 in undifferentiated (ESC), partially differentiated (NSC) and fully differentiated (preB) cell types persisted through the entire range of applied forces in our experiments.

### A progressive increase in metaphase chromosome stiffness can be modelled by local crosslinking

It is not immediately obvious what the underlying cause of the changes in chromosome biomechanical properties across different cell types can be. A plausible explanation for the increased metaphase chromosome stiffness in differentiated cells might be that the underlying chromatin acquires more stable, complex structures as cells become progressively more specialised. To try and model this, we examined the impact of experimentally inducing local crosslinking of chromatin/DNA using EGS. Specifically, we asked whether local crosslinking might be sufficient to alter the properties of ESC-derived metaphase chromosomes to more closely resemble equivalents in the more differentiated preB cells.

For this, we purified metaphase chromosome 3 from c10 ESCs as previously described and examined force extension characteristics of individual chromosomes after *in situ* treatment with crosslinker. As illustrated for a single chromosome in Figure 4c, local crosslinking with EGS (post, +EGS) resulted in reduced extension under force for the same individual chromosome when compared to before crosslinking (pre, −EGS). Crosslinking with EGS led to an average 15% decrease in the extension at 50 pN for each chromosome. Overall, this resulted in a mean extension of 1.97 ± 0.08 µm, similar to that for preB-derived chromosome 3. Crosslinking with EGS also led to overall higher force-dependent chromosome stiffness when compared to the untreated mouse ESC chromosome 3 (Figure 4d). At low forces (1 pN), the stiffness for EGS-treated chromosomes spanned the entire range from the low stiffness of ESC chromosomes to the higher stiffness of the preB chromosomes (Figure 4e). In contrast, under a high force (100 pN), the stiffness of EGS-treated ESC chromosomes significantly increased and more closely resembled the trend observed for NSC or preB-derived samples (Figure 4f). This trend is also observed in the force-dependent stiffening for each sample, given by the mean *γ* value (Figure 4g). Locally crosslinked ESC chromosomes exhibit the highest *γ* (0.87 ± 0.02), when compared to differentiated preB cells (0.84 ± 0.02), partially differentiated NSCs (0.78 ± 0.02), and undifferentiated ESCs (0.75 ± 0.02).

While we do not yet know the underlying cause behind how the mechanical behaviour of metaphase chromosomes is modulated at different stages of differentiation, the above results suggest that one possible explanation would be through changes in subchromosomal domain architecture (Figure 4h). The biomechanics of mitotic chromosomes arise from the force responses of these subchromosomal domains. Previous studies^15^ have suggested that chromosomes exhibit both fluid-like mechanical properties, which may arise from relative sliding between these domains, and solid-like properties, which may arise from reversible stretching of the domains themselves. Changes to this architecture, through local crosslinking, could reduce interdomain sliding, favouring intradomain stretching instead. Our data demonstrate that these crosslink-induced changes in architecture result in stiffer, more compact chromosomes with stronger force-dependent stiffening. It is likely that similar changes in subchromosomal architecture also underlie some of the biomechanical trends we observe across cell types at different stages of differentiation (Figure 4i).

## Discussion

Mammalian chromosomes are highly organised polymers that comprise nucleosomes (with linker DNA), accessory proteins, and RNA, and are subdivided into discrete compartments reflecting activity, topological interactions, repeat-DNA content, and modified histones. Entry into mitosis is a precisely choreographed process where increases in phosphorylation, mediated by dedicated kinases, alter key chromosome-associated proteins and lead to rapid eviction of many DNA-associated factors, loss of architectural features such as TADs, and physical compaction. While some of the mediators of this compaction (such as SMCs) are known^30–33^, the processes that shape mitotic chromosome conformation remain elusive.

In this study, we have examined the properties of defined mouse mitotic chromosomes, purified by flow cytometry from different cell types. From biomechanical stretching experiments using optical tweezers, we have shown that mouse chromosomes display non-linear stiffening behaviours under increasing force, consistent with reports from other groups studying human mitotic chromosomes^16^. Our data, however, suggest that mouse metaphase chromosomes are substantially less stiff (by nearly an order of magnitude) than those isolated from human U2OS cells. While this could be related to their structure, it is also possible that this reflects differences in experimental approach. For example, our studies were performed using stable endogenously generated BirA-TRF1 tags; furthermore, we examined chromosomes generated from diploid, non-malignant cell lines. Mouse chromosomes also have unique features including their acrocentric shape and repetitive DNA sequences contained within minor and major satellites contributing to centromere function, which fundamentally differ from those in humans in terms of monomer length, organisation, and location^22,34,35^. Whatever the underlying cause, we show that these mechanical properties are characteristic of both unsorted and unstained mouse chromosomes as well as sorted specific chromosomes. Therefore, we rule out any possibility that the reduced stiffness we observe is due to the purification procedure.

By studying purified individual metaphase chromosomes, we minimise heterogeneity arising from a mixed population of different chromosomes from our experiments. This reveals three aspects of chromosome biomechanical behaviour that were previously inaccessible. First, different chromosome types exhibit distinct chromosome-specific biomechanical properties, with mouse ESC chromosome 19 having higher stiffness than chromosome 3. However, despite these differences in stiffness, both ESC chromosomes share a conserved force-dependent stiffening behaviour. Second, the biomechanical properties of metaphase chromosomes vary based on the cell differentiation stage. A direct comparison of the properties of equivalent chromosomes derived from different cell types show that both the stiffness and force-dependent stiffening of chromosomes increase in more differentiated cells. Thirdly, even purified specific chromosomes can exhibit a wide range of heterogeneity in their mechanical response. By analysing more than 35 ‘identical’ chromosomes from each cell type, we show that chromosomal heterogeneity varies among the different mitotic chromosome preparations. Individual mitotic chromosomes isolated from mouse preB cells showed more similar extension profiles than mitotic chromosomes isolated from ESCs.

One explanation for the differences in variation between cell types could be that chromatin (and particularly heterochromatin) organisation is quite unusual in ESCs and radically changes as cells differentiate. This claim is supported by data from a range of orthogonal methods. For example, correlative electron spectroscopic imaging has been used to show that heterochromatin fibres are more dispersed in pluripotent cells than differentiated cells^36,37^ and that the unusually high levels of transcription of non-coding major satellite repeats typically seen in mouse ESCs, not only regulate the biophysical properties of constitutive heterochromatin, but are required to preserve genome stability through cell division^35^. An alternative but equally plausible explanation is that variation in mechanical properties is a consequence of the unusual cell cycle structure of mouse ESCs^38–40^. Mouse ESCs divide rapidly and have particularly short gap phases, so that at any time there is a high proportion of cells undergoing DNA replication (S-phase) or mitosis (M-phase). During synchronisation and arrest, it is possible that we sample a small number of cells entering mitosis (*i.e.* in prophase) in ESCs that are not represented in the slower dividing NSC or preB cell samples. Future studies will be needed to resolve this apparently heterogenous behaviours of sorted ESC-derived mitotic chromosomes. Nonetheless, the relatively wide extension distributions we observe for all cell types (and for different chromosomes from the same cell type) suggest that it is not possible to determine the identity of a specific chromosome from a mixed chromosomal pool by its force-extension behaviour alone, making purification techniques, like the advanced flow cytometry that we employ here, critical for investigating precise chromosome-specific biomechanics.

The progressive epigenetic changes that accompany differentiation are known to impact genome architecture and chromosome organisation. Currently our knowledge of how these changes alter chromosome folding is derived mostly from interphase studies using chromatin-based analysis combined with dimensional contact or DNA proximity assays. Species-specific and cell type-specific chromosome conformations^41^ may be relevant for chromosome compaction in mitosis^42^ as well as age-related and cell cycle-related changes in chromosome stiffness^43^. Heterochromatin features can impact mitotic chromosome structural folding and compaction^24,44^. Adding to these previous works, our data demonstrate that the progressive chromatinisation of the genome accompanying differentiation also alters the global physical and biomechanical properties of chromosomes in mitosis. Increased stiffness and force-dependent stiffening are hallmarks of more highly crosslinked polymer networks^45^, suggesting the presence of increasingly interconnected chromatin networks within the mitotic chromosomes of more differentiated cells.

Early micromanipulation experiments using isolated human mitotic chromosomes have provided evidence in support of a chromatin-network model^14^ where the chromatin fibre inside each chromatid is attached to itself by chromatin-chromatin crosslinks. In enzyme microspray experiments, Marko and colleagues reported that exposure of individual metaphase chromosomes to trypsin or proteinase K results in lengthening and thickening, with a progressive reduction in the doubling force. In contrast, exposure to restriction nucleases (4- base cutters that generate blunt ends), resulted in loss of chromosome elasticity and eventual chromosome dissolution. These data support the idea that chromatin crosslinks are a key load bearing element within mammalian mitotic chromosomes. A more recent study has shown that chromosomes can undergo phase transitions between loosely and more compact sub-chromosomal architectures controlled by noncovalent crosslinks^46^. Such non-covalent crosslinks could be mediated by electrostatic interactions between DNA phosphates and histone tails, which are known to be substantially modified during cell differentiation. An increase in such non-covalent crosslinks with differentiation stage can explain our observation that chromosomes from more differentiated cells are more compact and stiffer, providing a molecular model for how differentiation-related changes in chromatin chemistry can affect large-scale chromosome architecture and biomechanics. Such a model is further consistent with our own results that crosslinked chromosomes are stiffer than their non-crosslinked counterparts.

Flow cytometry-based separation of individual metaphase chromosomes has previously been used for detailed human karyotyping^47^, generating chromosome-specific probes^48^, as well as experimentally to identify potential mitotic bookmarking factors^23^, to interrogate CENPA structures at human centromeres and neocentromeres^49^ and in building new pipelines to engineer synthetic human chromosomes^50^. Here we describe its use in isolating genetically identical metaphase chromosomes from different cell types and differentiation stages for biomechanical investigation. This study provides a foundational cornerstone to investigate how individual metaphase chromosomes behave in different cell types, by comparing equivalents. Separation of individual mitotic chromosomes into their individual subtypes will be critical in accurately representing the mechanical impacts of changes to chromatin, for example by degradation or cleavage of remodelling factors, or SMCs. Undoubtedly, this will unlock a range of experiments to determine how epigenetic modifications^51,52^ can alter the biomechanics of metaphase chromosomes. Approaches of this sort enable us to move away from generic descriptions towards a clearer view of how biomechanical changes to chromosomes occur in development.

## Acknowledgements

We thank the MRC LMS Flow Cytometry Facility, the MRC LMS Microscopy facility, the Micron Bioimaging Facility (Department of Biochemistry, University of Oxford) and the Don Mason Facility of Flow Cytometry (Sir William Dunn School of Pathology, University of Oxford) for technical support. We thank Martin Houlard, Polina Maykova, William Allen and Adam Cawte for assistance with cloning constructs. This work was funded by the Medical Research Council UK (MC-PC-22015 awarded to A.G.F. and MC-A658-5TY10 awarded to D.S.R.). S.P. was supported by an MRC LMS Business Engagement Fund award (LMSBEF-GTEE-2024).

## Author contributions

A.G.F. and D.R. conceived the study and supervised the work. A.D., K.E.B., S.C., and M.E.K. generated and validated BirA-TRF1-expressing cell lines, with additional validation by M.L.A. and D.H.G. K.E.B. generated metaphase spreads and performed FISH experiments; C.W. performed size measurement analysis. A.D. extracted mitotic chromosomes, with assistance from K.E.B. and S.C.; B.P. isolated mitotic chromosomes by flow cytometry, with assistance from D.H.G. S.P. and K.K.R. performed optical tweezer experiments and formulated the single-molecule analysis pipeline. A.D., K.K.R., K.E.B., and S.P. prepared figures; A.G.F., A.D., S.P.,

K.K.R. and D.R. wrote the manuscript with input from all authors.

## Data availability statement

The associated data will be made freely available via a Zenodo repository at the time of publication.

## Methods

### Cell culture

E14Tg2a mouse ESCs (WT and BirA-TRF1 knock-in) were cultured on 0.1% gelatin in KnockOut DMEM medium (Gibco) supplemented with 10% FCS, non-essential amino acids, L-glutamine, penicillin/streptomycin, β-mercaptoethanol (all Gibco) and leukaemia-inhibitory factor (produced in-house).

For differentiating mouse ESCs into NSCs^29^, 0.45×10^6^ ESCs were seeded into a gelatin-coated T25 flask and cultured in N2B27 medium (1:1 mixture of DMEM/F-12 and Neurobasal medium, supplemented with 1× N2 (17502048), 1× B27 (without vitamin A, 12587010), 1 mM L-glutamine, 100 μM β-mercaptoethanol, antibiotics and non-essential amino acids (all Gibco). On day 7 of differentiation, cells were detached using TrypLE Express (Gibco), and 3×10^6^ cells were plated onto a 90 mm bacterial petri dish in N2B27 medium supplemented with 10 ng/ml EGF and FGF-2 (PeproTech, 315-09 and 100-18B). On day 10, cell clusters were collected by mild centrifugation (100 *g* for 2 min) and plated onto gelatin-coated culture dishes in N2B27 medium containing EGF and FGF-2. Once NPC outgrowths reached confluency, cells were detached with TrypLE Express and passaged at a 1:3 ratio onto fresh gelatin-coated dishes. After 2-3 passages, NSCs were transferred to culture dishes coated with laminin (Sigma) to support continued proliferation and were maintained in N2B27 medium supplemented with EGF and FGF-2.

Previously derived Abelson-transformed WT mouse preB cells^23,53^ were grown in IMDM medium supplemented with 12% FCS, 2 mM L-glutamine, 2X non-essential amino acids, penicillin/streptomycin and β-mercaptoethanol. All cells were cultured at 37 °C with 5% CO2. For biotin incorporation, biotin (Metasystems, B4501) was added to the culture medium for 24 h at a final concentration of 50 µM.

### CRISPR/Cas9-mediated *BirA-TRF1* knock-in

Guide RNA oligos targeting the *Rosa26* or TIGRE locus (Table 1) were annealed, phosphorylated (T4 Polynucleotide Kinase, NEB, M0201S) and assembled by golden gate assembly into the pU6-(BbsI)_CBh-Cas9-T2A-mCherry plasmid^54^ (gift from Ralf Kuehn, Addgene #64324) using BbsI-HF (NEB, R3539S) and T4 DNA ligase (NEB, M0202S). The *BirA-TRF1* transgene was amplified from a plasmid from R. J. O’ Sullivan^28^ (provided by Simon Boulton). A *BirA-TRF1* repair vector (Supplementary File 1) targeting the *Rosa26* locus was assembled using NEBuilder HiFi DNA Assembly Master Mix (NEB, E5520S) and a plasmid containing *Rosa26* homology arms (gift from the Brockdorff lab). A repair vector targeting the TIGRE locus was gifted by the Srinivasan lab (Supplementary File 2). Constructed plasmids were transformed into DH5α bacteria, purified and validated by sequencing (Source BioScience or Azenta Life Sciences).

Mouse ESCs or preB cells were co-transfected using Lipofectamine 3000 (Invitrogen, L3000008) with the sgRNA/Cas9 and BirA-TRF1 donor plasmids corresponding to the *Rosa26* or TIGRE loci respectively. The following day, mCherry-positive cells were bulk-sorted by flow cytometry and expanded for one week, before single-cell sorting into 96-well plates to derive clonal cell lines (without puromycin-based selection). Clones were screened and verified by PCR using the primer pairs indicated in Table 1. Heterozygous knock-in (KI) clones c10 (ESCs) and 2.12 (preB cells) were selected at random for downstream experiments.

**Table 1.**
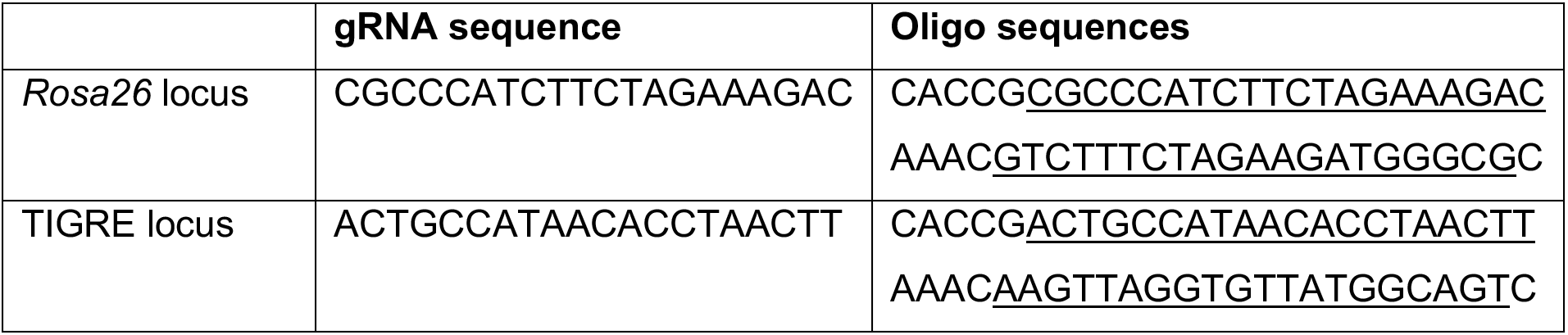
Guide RNA sequences.

**Table 2.**
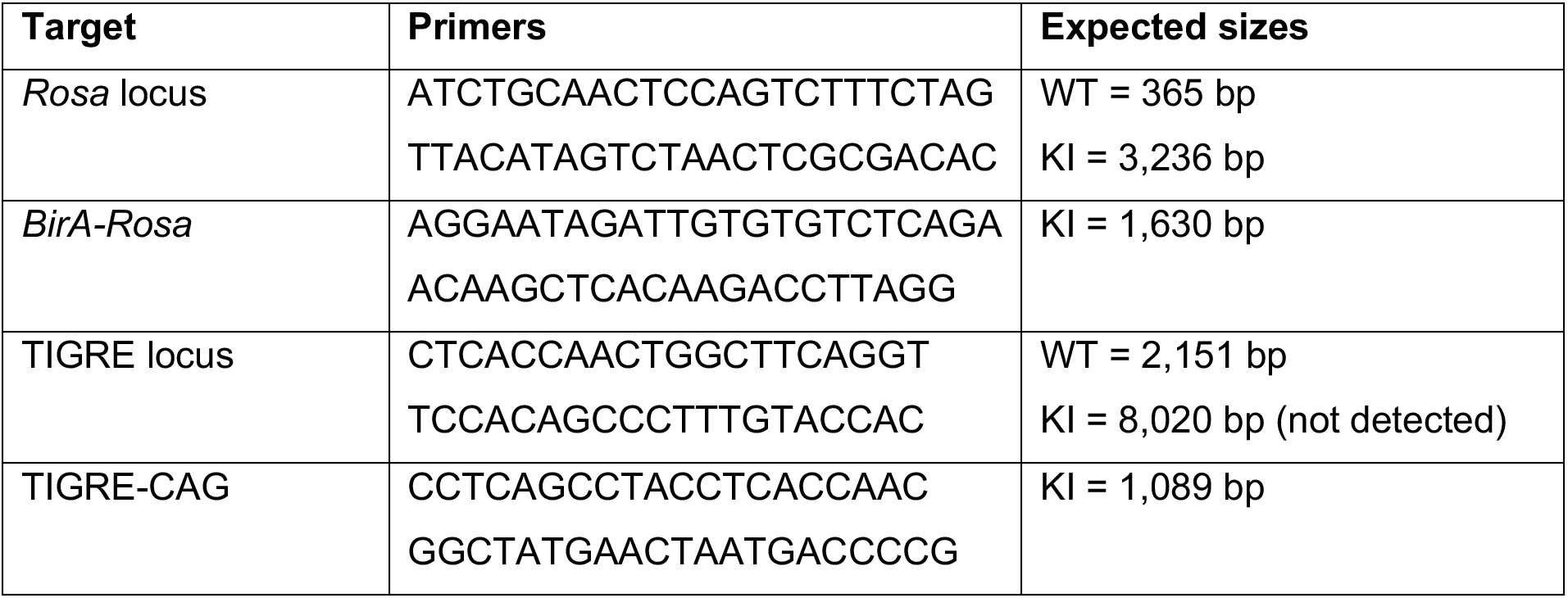
PCR primers for knock-in screening and validation.

### Immunofluorescence

For biotin detection (all steps at room temperature), cells were fixed in 4% formaldehyde (Thermo Scientific, 28906) for 10 min, washed twice in PBS, permeabilised with 0.5% Triton X-100 (Sigma, T-8787) for 10 min, washed twice in PBS and blocked with 5% FCS for 30 min. Cells were incubated for 1 h with streptavidin-Alexa Fluor 488 (Invitrogen, S11223) diluted 1:200 in blocking solution, washed with PBS, and mounted in Vectashield containing DAPI (Vector Laboratories, H-1200). Imaging was performed with a Leica STELLARIS 5 confocal microscope (LAS X software, v4.9.0) or an Olympus ScanR spinning disk confocal microscope equipped with a UPLXAPO 100× oil-immersion objective (Olympus cellSens Dimension software, v4.4). Representative images were processed in Fiji (1.54p)^55^.

For antibody staining (all steps at room temperature), cells were fixed in 2% paraformaldehyde (Fluka, 76240) for 20 min, washed thrice with PBS, permeabilised with 0.4% Triton X-100 for 5 min, washed twice with PBS and once in wash buffer (PBS with 0.2% BSA fraction V and 0.05% Tween-20 (Sigma, P-9416)), before blocking (10% FCS, 2.5% BSA Fraction V, 0.05% Tween-20 in PBS) for 30 min. Primary antibodies to OCT4 (Abcam, ab19857) and Nestin (Abcam, ab316018) were diluted 1:200 in blocking solution and incubated with cells for 1.5 h. Cells were washed thrice with wash buffer and incubated 30 min with donkey anti-rabbit secondary antibody (Alexa Fluor 488, Invitrogen, A-21206) diluted 1:400 in blocking solution. After washing twice in wash buffer and once in PBS, cells were mounted in DAPI-containing Vectashield. Images were collected with a Leica STELLARIS 5 confocal microscope and LAS X software (v4.9.0), and representative images were processed in Fiji (1.54p)^55^.

### Metaphase spreads and chromosome size measurements

Mouse ESCs were arrested with 0.1 μg/ml demecolcine (Sigma-Aldrich, D1925) for 2 h. Arrested cells were resuspended in hypotonic solution (75 mM KCl, 10 mM MgSO_4_, pH_RT_=8.0) at 37 °C for 25 min, before pelleting at 500 *g* for 8 min. Pellets were resuspended in residual supernatant, fixed in 75% methanol:25% acetic acid and stored at -20 °C. Pellets were washed three times in fixative (500 *g*, 8 min) and resuspended to a pale grey appearance in a small volume of fixative. A 20 μl drop of 45% acetic acid was placed on a glass Twinfrost slide and 23 μl of sample was dropped directly on top, tilting to spread chromosomes. Slides were air-dried at room temperature before further processing.

Mouse chromosome paints for chromosomes 3 and 19 (MetaSystems Probes, D- 1403-050-OR and D-1419-050-FI) were hybridised to samples according to the manufacturer’s protocol before mounting in Vectashield containing DAPI (Vector Laboratories). Images were collected with a Leica STELLARIS 5 confocal microscope using LAS X software (v4.9.0). Images were segmented with Cellpose (v3.1.1.2)^56^, using the ’cyto3’ model with additional training using manual corrections to enhance segmentation within clumped spreads, and QuPath (v0.6.0)^57^ was used for chromosome classification and area measurements. Representative images were processed in Fiji (1.54p)^55^. GraphPad Prism (v11.0.1) was used for statistical analysis and preparing plots.

### Extraction, staining and flow cytometry-based purification of mitotic chromosomes

Mitotic chromosomes were isolated using flow cytometry as previously described^23^. Cells were first arrested for 5 h with 0.1 μg/ml demecolcine (Sigma-Aldrich, D1925). For mouse ESCs and NSCs, mitotic cells from ten 100 mm dishes were collected by harvesting the medium and any detached cells following 1-2 min room temperature TrypLE Express treatment. For mouse preB cells, approximately 10^8^ demecolcine-treated cells were used. Arrested cells were pelleted and resuspended in 10 ml of hypotonic solution (75 mM KCl, 10 mM MgSO4, 0.2 mM spermine tetrahydrochloride (Sigma-Aldrich, S2876), 0.5 mM spermidine trihydrochloride (Sigma-Aldrich, S2501), pH_RT_=8.0) and incubated at room temperature for 20 min. Cells were resuspended in 1–3 ml of polyamine (PA) buffer (80 mM KCl, 15 mM Tris-HCl, 2 mM EDTA, 0.5 mM EGTA, 3 mM DTT, 0.25% Triton X-100, 0.2 mM spermine tetrahydrochloride, 0.5 mM spermidine trihydrochloride, pH_RT_=7.7) and incubated for 15 min on ice. Samples were vortexed for 30 s (maximum speed) and passed several times through a 21-gauge needle. Intact cells and debris were removed by centrifugation (200 *g* for 2 min) and filtering (20 μm CellTrics filter, Sysmex, 04-004-2325). Chromosomes were stored at 4 °C overnight before staining for 1 h on ice with Hoechst 33258 (5 μg/ml final, Sigma-Aldrich, 94403), chromomycin A3 (25 μg/ml final, Sigma-Aldrich, C2659), MgSO_4_ (10 mM final) and, for centromere imaging experiments, Alexa Fluor 647-conjugated anti-CENP-A antibody (1:200, Cell Signaling Technology, #2048, with custom fluorophore conjugation). Sodium citrate (10 mM final) and sodium sulphite (25 mM final) were then added, and samples were incubated on ice for a further 1 h before flow purification. Stained chromosomes were purified on a Becton Dickinson Influx (nozzle tip = 70 μm, drop-drive frequency = 96 kHz, sheath pressure = 448 kPa, BD FACS Sortware v1.2.0.142). Forward scatter was measured with a 488 nm laser (Coherent Sapphire, 200 mW). Hoechst 33258 was measured with either a 355 nm air-cooled laser (Spectra Physics Vanguard, 350 mW) or a 355 nm water-cooled laser (Coherent Genesis, 250 mW), coupled to 400 nm (long-pass)/500 nm (short-pass) filters. Chromomycin A3 was measured with a water-cooled 460 nm laser (Coherent Genesis, 500 mW), coupled to 500 nm (long-pass)/600 nm (short-pass) filters.

To assess sort purity, flow-isolated chromosomes were cytocentrifuged (Cytospin3, Shandon) for 10 min at 1,000 rpm (∼163 *g*) onto Polysine slides (VWR, 631-0107) and hybridised with paints for chromosomes 3 and 19 (MetaSystems Probes, D-1403-050-OR and D-1419-050-FI) according to the manufacturer’s protocol. Samples were mounted in Vectashield containing DAPI (Vector Laboratories) and images were collected on a Leica STELLARIS 5 confocal microscope using LAS X software (v4.9.0). Chromosome paint- positive and negative chromosomes were counted manually and representative images for display were processed in Fiji (1.54p)^55^.

### Reverse transcription quantitative PCR

Total RNA was isolated from 1×10⁷ cells using TRIzol reagent (Invitrogen, 15596026), and first-strand cDNA was synthesised from 2 µg of total RNA using random primers and SuperScript III reverse transcriptase (Invitrogen, 18080093) according to the manufacturer’s instructions. Quantitative PCR was carried out on a CFX96 Real-Time System (Bio-Rad; CFX Manager v3.1), using Applied Biosystems SYBR Green Universal Master Mix (Thermo Fisher Scientific, 10607705) and primers listed in Table 3, with each reaction pipetted in technical triplicate. Gene expression was calculated relative to the average Ct values of *Gapdh* and *β- actin (Actb)*. Plots were prepared in GraphPad Prism (v11.0.1).

**Table 3.**
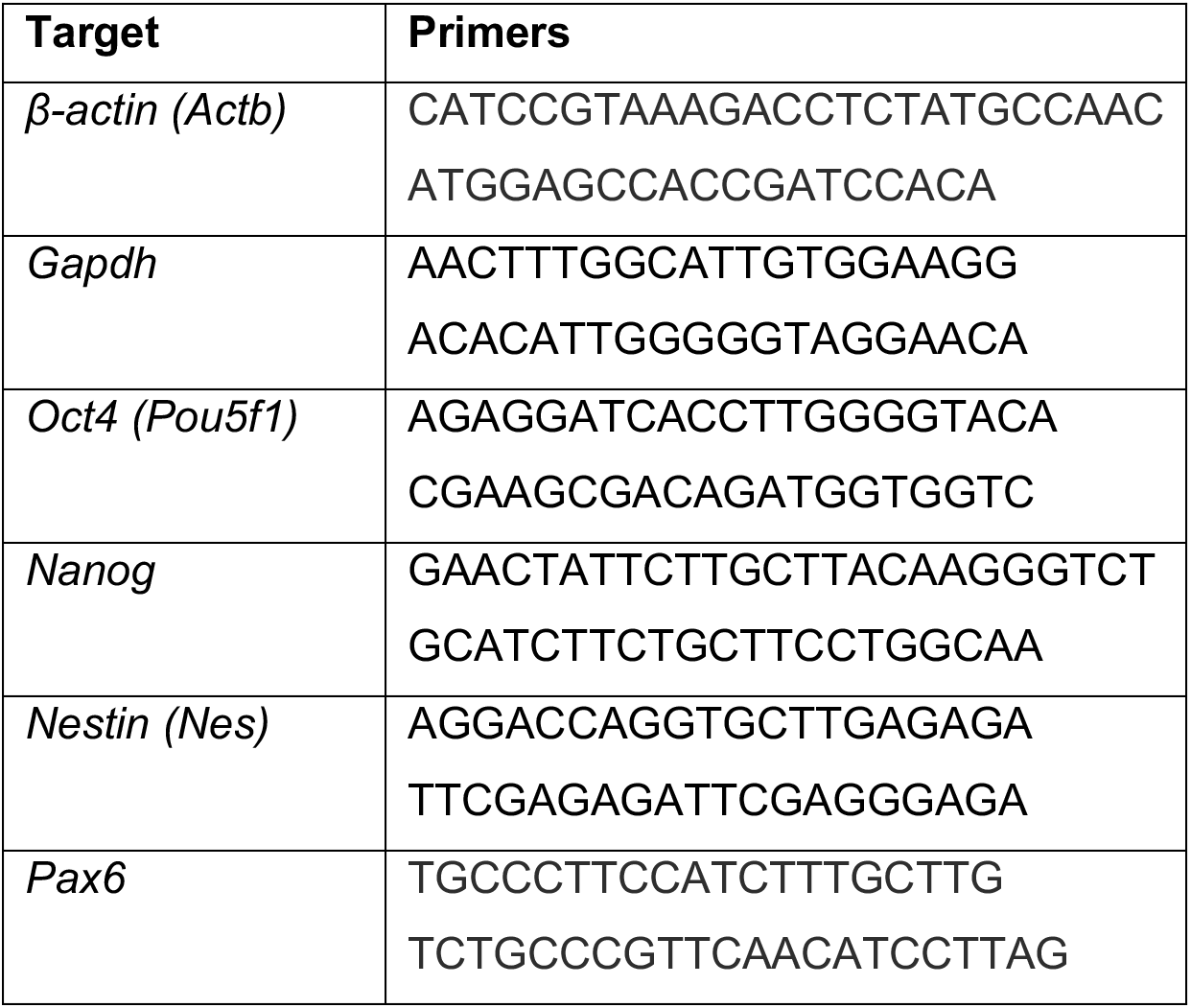
qRT-PCR primers.

### Optical tweezers experiments

Single-chromosome experiments were performed using a C-Trap (LUMICKS) integrating optical tweezers, confocal fluorescence microscopy, and microfluidics. Before each experiment, the five-channel microfluidic flow cell was passivated by incubation with 0.05% (w/v) casein in TP50 buffer (50 mM Tris-Cl pH_RT_=7.5, 50 mM KCl) for 1 h followed by extensive rinsing of channel 1 with TP50 buffer and the rest of the channels with PA buffer for 1 h. At the start of the experiment, the following were introduced to the microfluidic channels: (i) 0.005% (w/v) streptavidin-coated polystyrene beads (Spherotech, 4.35 µm and 2.17 µm for chromosomes 3 and 19, respectively) in TP50 in channel 1; (ii) PA buffer in channels 2 and 3; (iii) specific flow-sorted biotinylated chromosomes in PA buffer (at a concentration of ∼10^6^ chromosomes per 1 ml) in channel 4.

The optical traps were operated at a trap stiffness of 0.3–0.5 pN/nm. Individual chromosomes were tethered between two optically trapped beads as previously described^16^. Briefly, one optically trapped bead was first brought in proximity of an individual chromosome in channel 4, allowing the biotinylated telomere to attach to the streptavidin-coated bead. Following attachment, the chromosome-bound bead, along with the second optically trapped bead, were transferred to channel 3. The chromosome was briefly stretched under controlled buffer flow to confirm successful attachment. The second bead was brought near the other end of the flow-stretched chromosome to form a second attachment.

After the formation of the tether, multiple force-extension measurements of the individual chromosome were carried out in PA buffer by moving one optical trap at a constant speed (20–100 nm/s). Bright-field images of the force-extension measurements were recorded. Chromosomes that displayed visible damage, aggregation, or unravelling were excluded from further analysis. Fluorescent images of the individual tethered chromosomes were captured by exciting the chromosome-bound chromomycin A3 with a 488 nm laser (laser power 8 µW). For chromosomes with labelled CENP-A, the Alexa Fluor 647-conjugated CENP-A specific antibody was detected using a 638 nm laser (laser power 2 µW).

### *In situ* chromosome crosslinking

For crosslinking experiments, a saturating concentration of ethylene glycol bis(succinimidyl succinate) (EGS) in Crosslinking Buffer (50 mM HEPES pH=7.5, 80 mM KCl, 10 mM MgCl_2_, 2 mM EDTA, 0.5 mM EGTA, 3 mM DTT, 0.25% Triton X-100) was introduced to channel 5 of the C-trap microfluidic chamber. An individual ESC chromosome 3 was first captured between optically trapped beads and stretched to 100 pN in PA buffer. The chromosome was then transferred to channel 5 and incubated in EGS for 1–2 min. The chromosome was then returned to the PA buffer channel, which quenched the remaining EGS. Force-extension measurements were performed under the same conditions as untreated chromosomes.

### Single-molecule data analysis

All single-molecule datasets represent data collected from at least 3 independent chromosome purifications. All images were analysed in Fiji (version 1.54t). The force-extension data were analysed using custom Python scripts implemented in Jupyter Notebooks using the Pylake package (LUMICKS).

Chromosome extensions were extracted at a fixed force of 50 pN. For each force-extension curve (FEC), this was determined as the extension of the chromosome at the largest force just below 50 pN. Curves for which no data points were recorded above 50 pN were excluded from the analysis. For each distribution, the coefficient of variation (*CV*) was calculated as 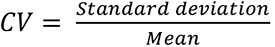

The stiffness (*K*) was calculated as

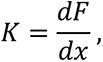

where *F* is the force and *x* is the extension. For each FEC, the force and extension were first smoothed using a uniform 1D filter over a window corresponding to 1/15^th^ of the total number of data points. The stiffness was then calculated for each point of the smoothed FEC using a numerical gradient. To reduce noise introduced by numerical differentiation, the calculated *K* values were subsequently smoothed using a uniform 1D filter with the same window as above. To determine the stiffnesses of the chromosomes at 1 pN and 100 pN, the smoothed stiffness-force data were interpolated onto a common force grid (0.1 pN).

The force-dependent stiffening behaviour of chromosomes was quantified using the stiffening exponent, *γ*, defined as

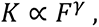

where *K* is the stiffness and *F* is the force. Rearranging the above, the local stiffening exponent for each force was calculated as

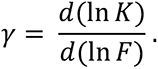

Values of *γ* outside the range of −10 to 10 were removed to exclude numerical artefacts. The local *γ* values thus obtained were then averaged to obtain a single representative *γ* for each FEC.

Statistical comparisons of chromosomal properties were performed using Welch’s unpaired t-test (with the Holm-Šídák correction for multiple tests wherever applicable), where *p* < 0.05 was considered statistically significant. The *CV* between two groups were compared using the asymptotic Feltz-Miller test for equality of *CV*s. The test statistic was evaluated against a χ^2^ distribution with one degree of freedom, with *p* < 0.05 considered statistically significant^58^.

---

**Supplementary Figure S1.**
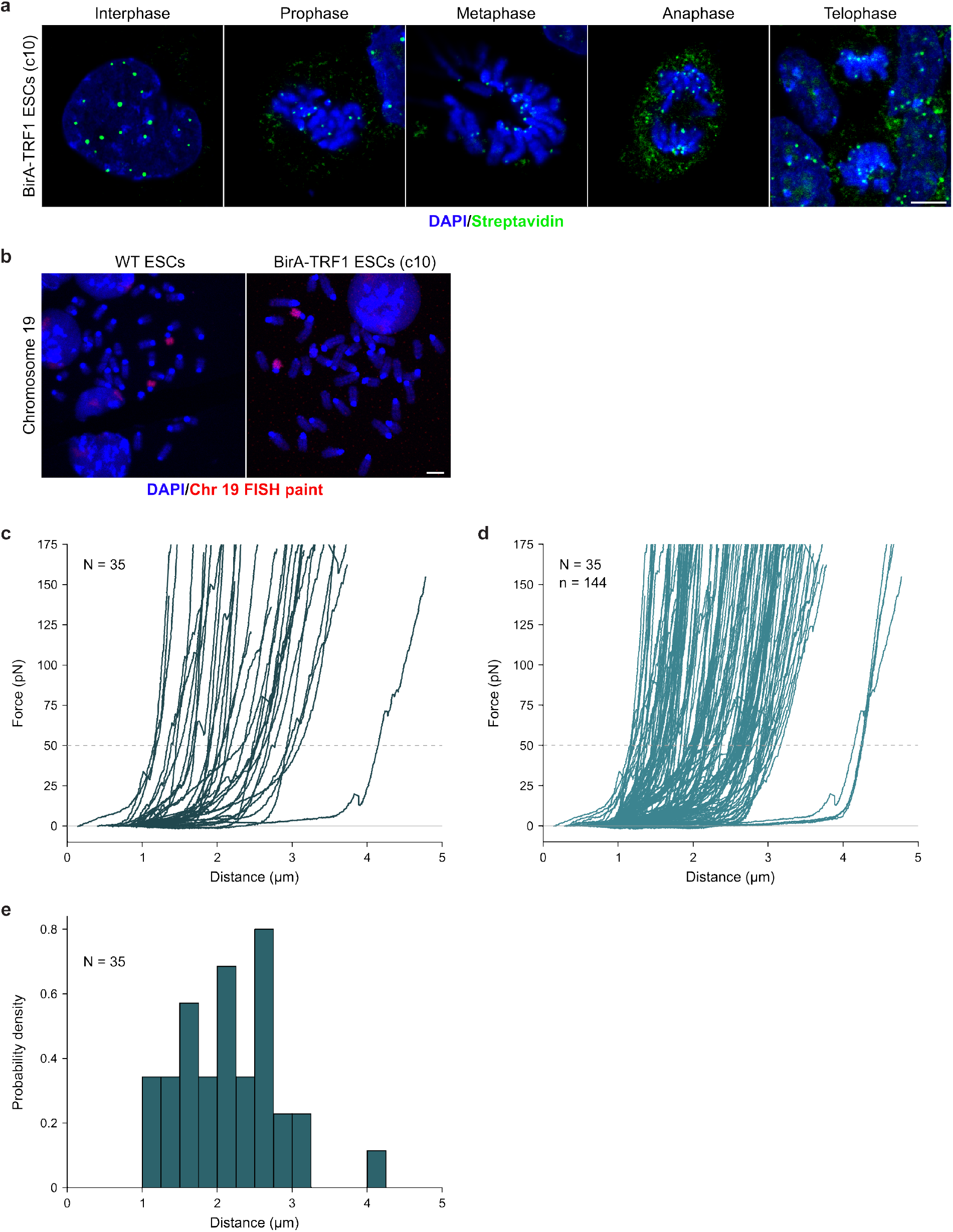
Characterisation of BirA-TRF1-expressing mouse ESC chromosomes. **a,** Detection of biotin incorporation at telomeres in BirA-TRF1-expressing c10 ESCs, using Alexa Fluor 488-labelled streptavidin. Single z-slices of representative cells from interphase and mitotic stages are shown. Scale bar = 5 µm. **b,** Representative metaphase spreads from parental WT and c10 (BirA-TRF1-expressing) ESCs, hybridised with chromosome 19 paint probes. **c,** Force-extension curves (FECs) for unstained and unsorted mouse ESC chromosomes, showing the first extensions (N = 35). **d,** FECs for unstained and unsorted mouse ESC chromosomes, showing all extensions (N = 35, n = 144). **e,** Histogram of unstained and unsorted mouse ESC chromosome extensions at 50 pN (N = 35). Extensions were calculated from the first FEC of each chromosome.

**Supplementary Figure S2.**
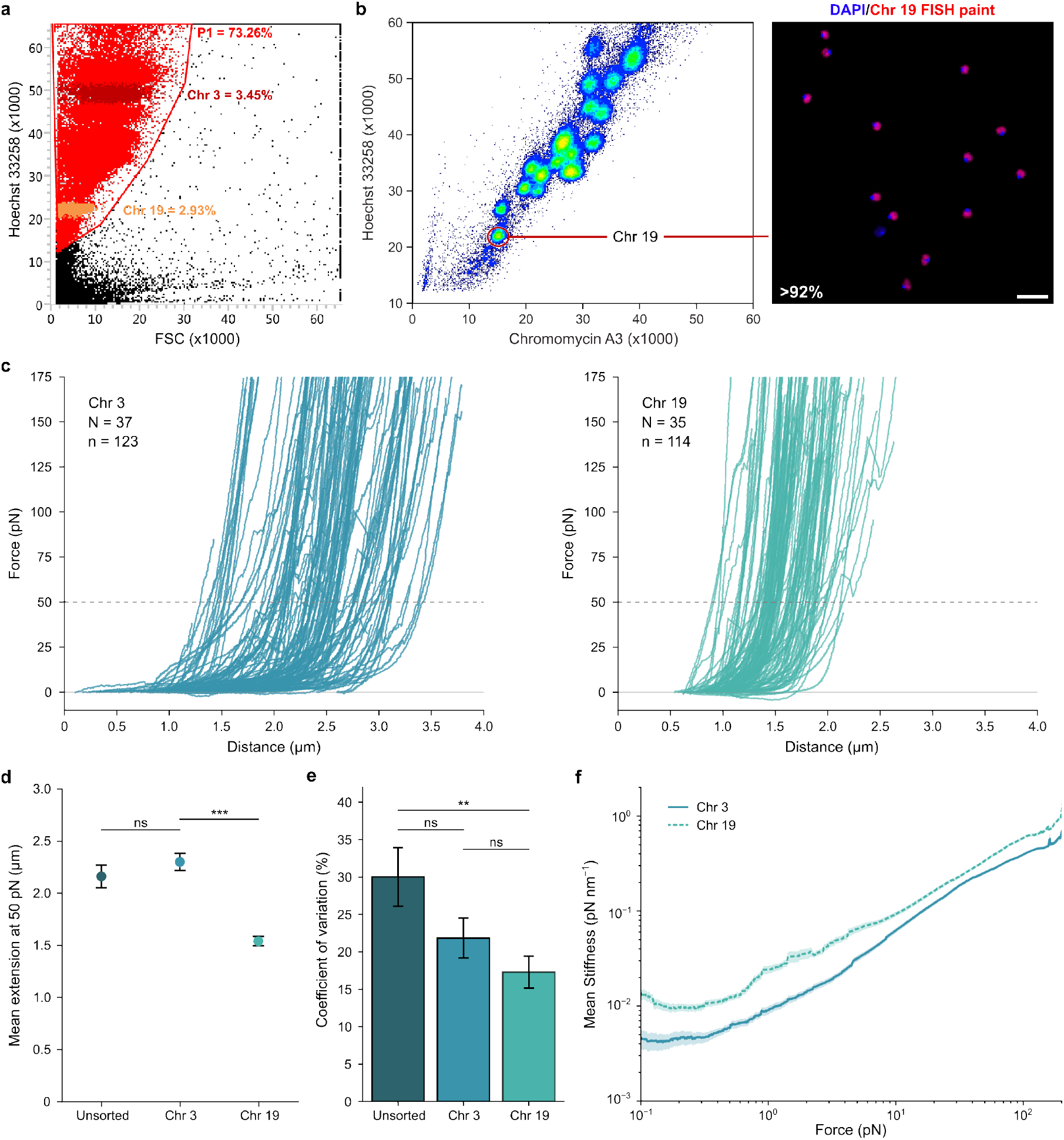
Biophysical properties of specific mouse ESC mitotic chromosomes. **a,** P1 gating strategy for flow cytometry-based purification of individual mitotic chromosomes from a representative experiment corresponding to the P1 events shown in Figure 2a and Supplementary Figure 2b. Percentages indicate the proportion of total events within the P1 gate and chromosome 3/19 events relative to the P1-gated population. **b,** Flow karyotype (left) of ESC mitotic chromosomes from Figure 2a, showing gating strategy for purification of chromosome 19. Purity was confirmed as >92% by hybridisation with chromosome 19 paint probes (right, N = 383). Scale bar = 10 µm. **c,** FECs for flow-purified c10 ESC chromosome 3 (left, N = 37, n = 123) and flow-purified c10 ESC chromosome 19(right, N = 35, n = 114). The first extensions of these samples correspond to Figures 2c and 2f, respectively. **d,** Mean extensions for unsorted ESC mitotic chromosomes (N = 35), flow-purified ESC chromosome 3 (N = 37), and flow-purified ESC chromosome 19 (N = 35) at 50 pN (Welch’s t-tests with Holm-Šídák correction, ns = not significant, *** = p_adj_ < 0.001). Error bars correspond to standard error of the mean (s.e.m.). **e,** Coefficients of variation for the extensions for unsorted ESC mitotic chromosomes (N = 35), flow-purified ESC chromosome 3 (N = 37), and flow-purified ESC chromosome 19 (N = 35) at 50 pN (asymptotic Feltz-Miller tests with Holm-Šídák correction, ** = p_adj_ < 0.01, ns = not significant). Error bars correspond to s.e.m. **f,** Force-stiffness curves for ESC chromosomes 3 (N = 37, n = 123) and 19 (N = 35, n = 114), shown on a log scale (corresponding to Figure 2h). Mean stiffness ± s.e.m. (shading) is shown. Extensions were calculated from the first FEC of each chromosome, while stiffnesses were calculated from all FECs.

**Supplementary Figure S3.**
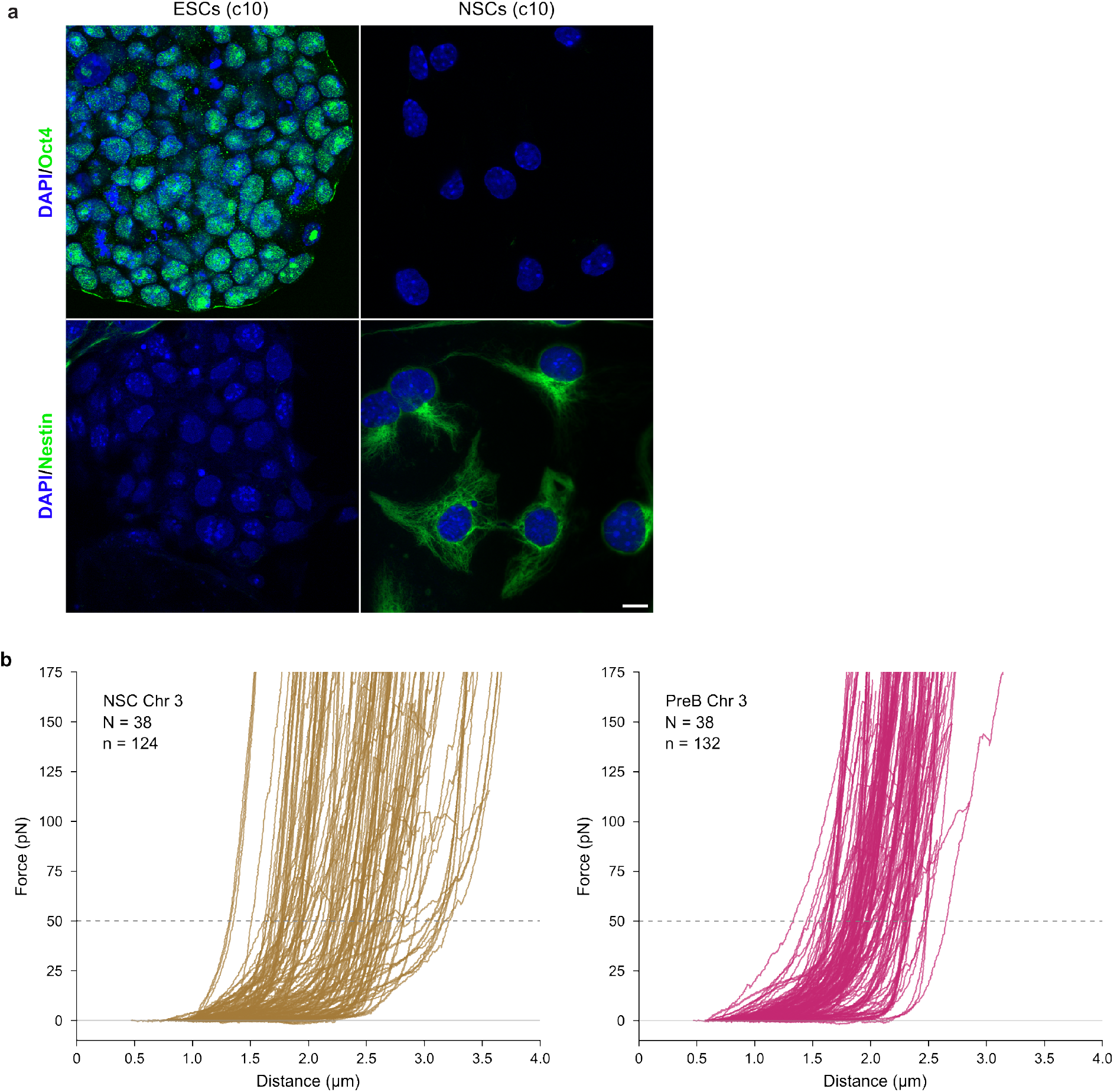
Biophysical analysis of mitotic chromosomes from differentiated cell lines. **a,** Antibody staining for Oct4 (upper) and Nestin (lower) in c10 ESCs (left) and after differentiation into self-renewing NSCs (right). Scale bar = 10 µm. **b,** FECs for flow-purified c10 NSC chromosome 3 (left, N = 38, n = 124) and flow-purified c2.12 preB chromosome 3 (right, N = 38, n = 132). The first extensions of these samples correspond to Figures 3c and 3h, respectively.

**Supplementary Figure S4.**
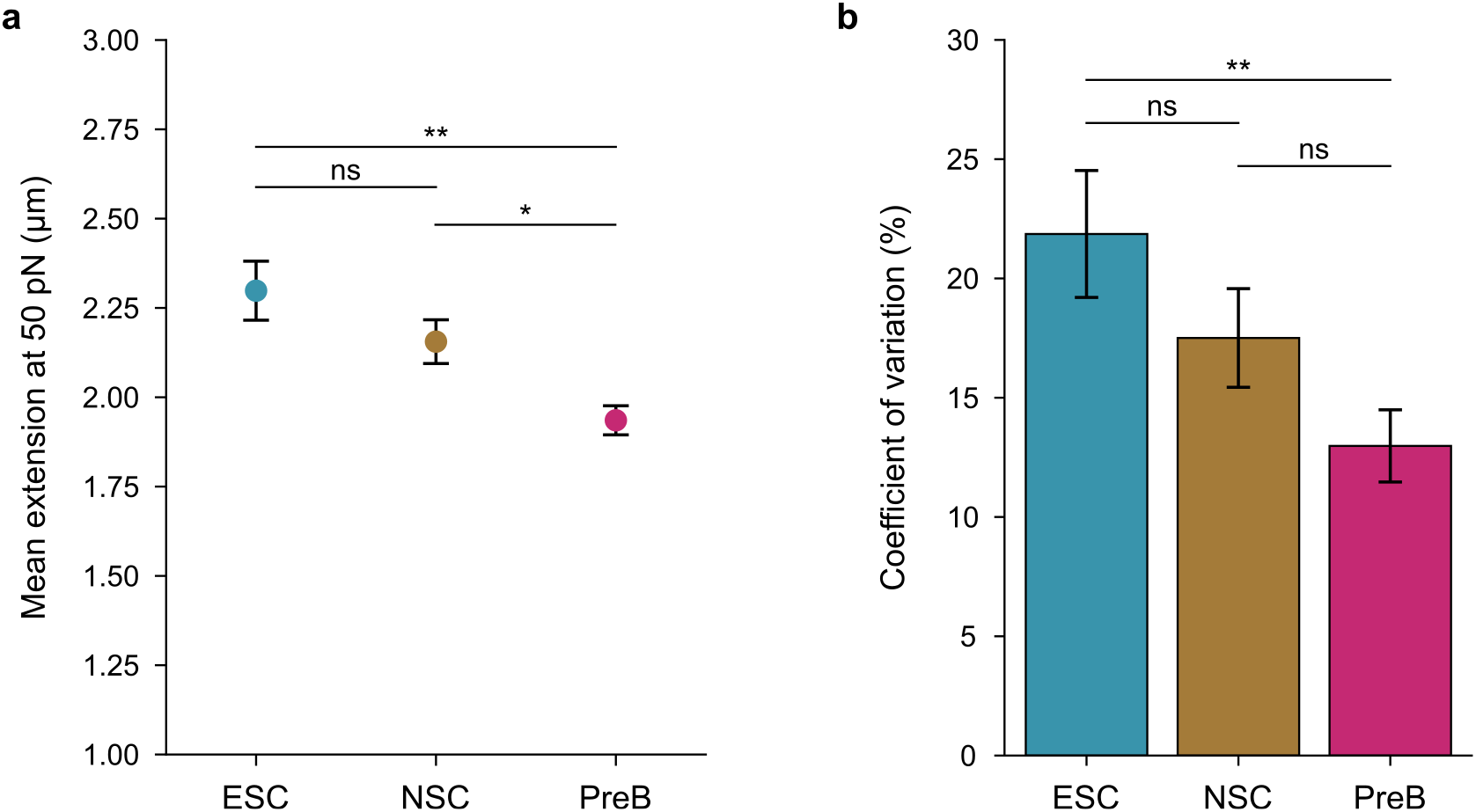
Biophysical properties of mitotic chromosomes from mouse ESC, NSC, and preB cells. **a,** Mean extensions for mitotic chromosome 3 from ESCs (N = 35), NSCs (N = 38), and preB cells (N = 38) at 50 pN (Welch’s t-tests with Holm-Šídák correction, ns = not significant, * = p_adj_ < 0.05, ** = p_adj_ < 0.01). Error bars correspond to s.e.m. **b,** Coefficients of variation for the extensions for mitotic chromosome 3 from ESCs (N = 35), NSCs (N = 38), and preB cells (N = 38) at 50 pN (asymptotic Feltz-Miller tests with Holm-Šídák correction, ** = p_adj_ < 0.01, ns = not significant). Error bars correspond to s.e.m.

